# shinyUMAP: an online tool for promoting understanding of single cell omics data visualization

**DOI:** 10.1101/2025.08.27.672621

**Authors:** Rohan Misra, Kevin O’Leary, Wenna Chen, Deyou Zheng

**Author notes:** Correspondence should be addressed to, Deyou Zheng, Ph.D.

## Abstract

Visualization is widely used to help explore and interpret high dimensional single cell (sc) omics data, such as scRNA-seq expression data. In particular, uniform manifold approximation and projection (UMAP) has become nearly ubiquitous in scientific publications that apply single cell omics technologies. Some experts have expressed concerns that the global cell-cell relationship, especially the spatial distances among cell clusters in a dataset, may not be faithfully depicted in a 2-dimensional (2D) UMAP. To help users to better appreciate this issue with their own data, we created an online server for the community to upload their single cell data and interactively make UMAPs with different hyper-parameters to witness how the distribution of cell clusters changes. The server thus can help promote proper usages of UMAP, especially to avoid the common pitfalls in misinterpretation of inter-cluster relationships in single cell studies.

**Availability and Implementation:** ShinyUMAP is freely available as an online Shiny server implemented in Python at https://scviewer.shinyapps.io/shinyUMAP/.

## Main Text

By resolving cellular heterogeneity, single cell technologies have become increasingly vital in the study of development, diseases, and cancers. These technologies undergo active development and novel applications emerge continuously. Among them, two mature and widely used methods are single cell RNA sequencing (scRNA-seq) for profiling transcriptomes ^1^ and single cell assay for transposase-accessible chromatin sequencing (scATAC-seq) for profiling chromatin accessibilities ^2^. The data scale, quality, and software for its analysis and integration have been significantly improved over the last few years, but visualization of the data (i.e. low-dimensional embedding of the cells in either 2D or 3D) remains relatively unchanged ^3^, with the early usage of t-distributed stochastic neighborhood embedding (t-SNE) ^4^,^5^ gradually replaced by uniform manifold approximation and projection (UMAP) ^6^,^7^. Other algorithms like Diffusion Map ^8^ and Potential of Heat-diffusion for Affinity-based Transition Embedding (PHATE) ^9^ also exist, but their usages are less prevalent ^10^.

It is generally accepted in the field that UMAP better preserves cell-cell relationship (or data structure) globally than t-SNE, but some disagree^11^,^12^. On the other hand, researchers have raised concerns that UMAPs are often mis-used in many single cell publications and some even suggested to avoid them entirely or limit their usage ^13, 14, 15, 11^. An appropriate consensus alternative, however, has not emerged. Essentially, embedding of cells in a UMAP (or t-SNE) is not a linear dimensional reduction but manifold approximation, with many parameters involved in the computing. While some prominent experts have consistently pointed out the limitations, many users in the wider community either are not fully aware of the computational details or choose to ignore them by using default setting in a software package. Furthermore, it is quite common to find authors describe two cell clusters (or types) in a UMAP as functionally more related or developmentally more connected simply because they appear closer than others in a UMAP. In reality, the spatial distance of cell clusters in a UMAP is affected by many tunable parameters in the UMAP computation, whereas the local structure (i.e., cells in the same cluster) is less influenced. Such parameters include “number of neighbors”, “minimal distance”, “number of components”, and others. We think the lack of interactive tools could be a reason why this issue has not been more widely appreciated. To fill this need, we decided to develop an online server that would enable users with less resources or computational skills to interactively and easily adjust parameters for UMAP computation, directly visualize their impacts, and save the results. Furthermore, we believe that researchers will better appreciate the impacts of parameters when the effects are illustrated with their own data, compared to theoretical explanations ^16^ or using prebuilt data (e.g., “Tensorflow Embedding Projector”^17^). Ultimately, such a tool would help bioinformaticians convey the information to experimental investigators who play critical roles in interpreting UMAPs.

We called this online tool shinyUMAP because it is built on a shiny server (**Figure 1**). We believe that its usage will shed light on the “blackbox” of UMAP generation. At its core, the server implements the UMAP function in the Scanpy package ^18^ and takes an AnnData input file (“h5ad” format) (**Figure 1A,B**). Once the input file is read, the server will plot the UMAP if it is included in the data object, colored by a metadata class. In cases where a dataset has not been processed in advance, shinyUMAP has an option to process the data using the standard Scanpy workflow, which is recommended by the scverse community.

**Figure 1.**
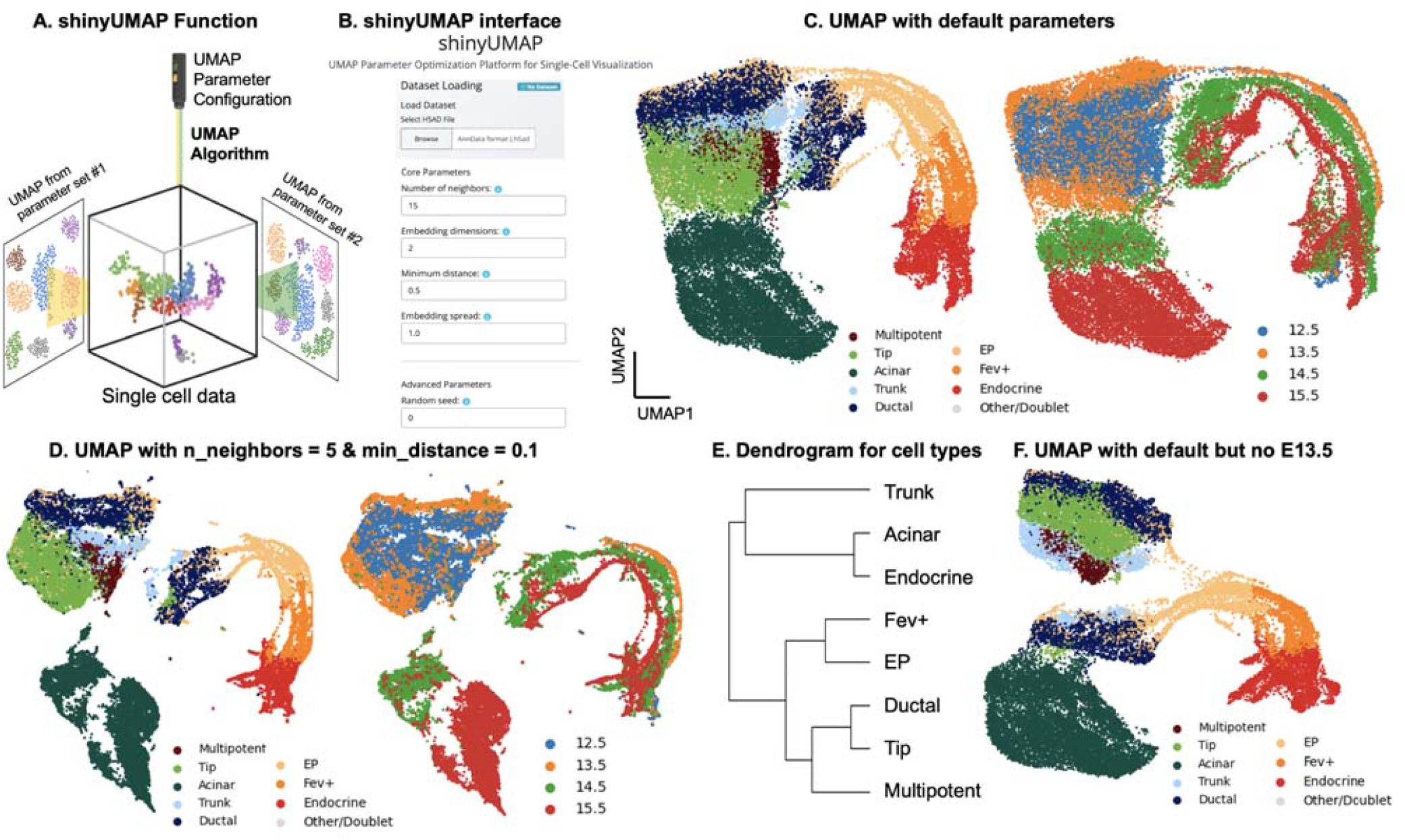
Usage of shinyUMAP. A) Cartoon indicating that the primary usage of shinyUMAP is to project users’ single cell data with different parameters (represented by yellow and green buttons of a flashlight). B) Key UMAP parameters in the shinyUMAP interface. C) UMAPs computed with default shinyUMAP parameters for a mouse pancreatic endocrinogenesis scRNA-seq dataset, colored by cell types (left) and embryonic days (right). D) UMAP after changing two key parameters, colored by cell types (left) and times (right). E) Dendrogram plotted by shinyUMAP to represent the gene expression similarity among the cell types. F) UMAP computed with default shinyUMAP parameters after cells in E13.5 were excluded.

To reduce memory usage and computational complexity, we prefer users submit data containing only the top variable features (i.e., highly variable genes in scRNA-seq or peaks in scATAC-seq vs all features) that are used for principal component analysis (PCA) and UMAP reduction. We also offer to randomly sample a smaller set of cells from input data if it contains a very large number of cells (e.g., > 50,000 cells). The users can then modify a set of parameters typically used in generating UMAPs such as “number of neighbors”, “minimal distance”, or “spread”, with a popup explaining the potential effect of each option. After users select and submit their parameters, a new UMAP will be computed and plotted, typically taking a few minutes. The new plots, UMAP coordinates, the corresponding parameters, and the updated AnnData object can be downloaded. These results can be used for downstream analysis, including single-cell dubious embedding detecting (scDEED) ^19^.

To illustrate the function of shinyUMAP, we applied it to a previously published scRNA-seq dataset for studying pancreatic endocrinogenesis by Bastidas-Ponce *et al*. ^20^. The dataset is from mouse embryos, containing 36,351 cells and 17,327 genes. **Figure 1C** shows the UMAP computed with default parameters in shinyUMAP (n_neighbors = 15, min_dist = 0.5), colored by the author defined clusters (or cell types; Multipotent, Tip, Acinar, Trunk, Ductal, EP, Fev+, Endocrine). The data were obtained from four developmental embryonic stages, E12.5 to E15.5 (**Figure 1C**, right), revealing the developmental lineage. Our UMAP matched the authors’ original (**Figure S1**), except the second dimension is flipped. We then changed the min_dist to 0.1 and n_neighbors to 5. The resulting UMAPs are shown in **Figure 1E** (by cell types on the left and by time on the right), with cells clustered more tightly and the appearance of small islands of cells as expected from the new parameters. A comparison of these UMAPs indicate that cells of the same type remain close to each other, but the relative positions of some clusters change and become more spread out. If a user treated the cluster distances as biologically meaningful, they would think that Tip and Acinar cells maintain developmental connections to the endocrine lineage (EP, Fev+, Endocrine) based on **Figure 1C**, consistent with the authors’ findings that tip cells contribute to endocrine progenitors during early development. However, this biological relevance would appear severed in **Figure 1D**, potentially leading researchers to conclude that tip and acinar lineages are completely separated from endocrine development. To help with this, shinyUMAP produces a dendrogram to show the gene expression similarity among cell types (or clusters) (**Figure 1E**). The dendrogram therefore provides information about potential functional relationships among clusters. Surprisingly, in this case, Acinar and endocrine (both derived from a common multipotent progenitor) have very similar expression profiles but are distinct in the UMAPs. To further demonstrate that cell type positions in an UMAP need to be interpreted with caution, we removed cells from one of the four time points (E13.5) and re-computed a UMAP with default parameters (**Figure 1F**). A comparison of the UMAPs in Figure 1C and 1F shows a major difference in Tip cell positioning. Without E13.5 cells, they lose connections to the Acinar cells. Thus, this example clearly illustrates how shinyUMAP can help users better understand how the UMAPs from their data are dependent on hyper-parameters. The appearance of a UMAP depends entirely on what data users have in hand (which could miss some cell types or developmental stages) and what UMAP parameters they select.

Additional examples using scRNA-seq or scATAC-seq data from several published datasets are included in the Supplement (**Figure S2-S5**) and further demonstrate that the influence of parameter choices on UMAPs affect some datasets more profound than others.

The shinyUMAP code is written in python and thus cannot be applied to objects from R workflow, such as Seurat directly. To address this, we provide a script for format conversion, but users can also use a recently developed anndataR package ^21^. Another limitation is that shinyUMAP cannot handle big datasets that exceed the memory allocated to use in the Shiny environment. To overcome this, we plan to develop a shiny-independent package that users can install and run locally.

## Acknowledgement

We thank our lab members for suggestion and extensive testing of the shinyUMAP server.

