## Supplemental Figures S1-S5 for "shinyUMAP: an online tool for promoting understanding of single cell omics data visualization"

**Supplemental Analysis and Figures S1-S5.**

**Figure S1. UMAPs computed by the original authors of the mouse pancreatic scRNA-seq data.** UMAPs generated by shinyUMAP using the coordinates in the AnnData object provided by the authors of the mouse pancreatic endocrinogenesis scRNA-seq data [1], colored by cell types (left) and embryonic (E) days (right).

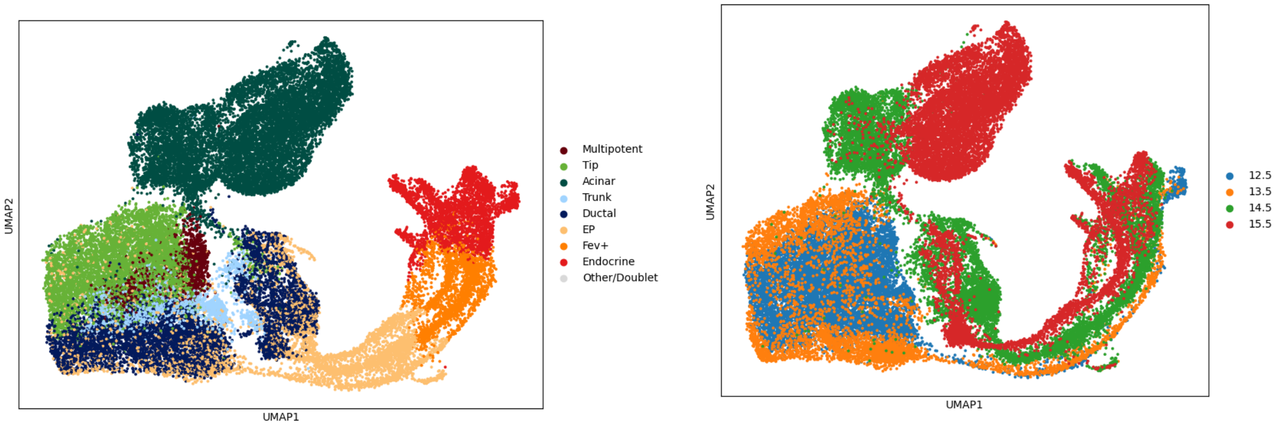


### I. Applying shinyUMAP to a human peripheral blood mononuclear cell (PBMC) dataset from patients with Coronavirus disease 2019 (COVID-19) and influenza

PBMC scRNA-seq data was obtained from a previous study of immune response following SARS-CoV-2 infection [2]. To reduce data size, we excluded cells from healthy controls, as described in our previous publication [3] , which resulted in 35,304 cells separated into 15 cell types/clusters defined in the original study. We uploaded the dataset to shinyUMAP and generated a UMAP with default parameters, colored by cell types (**Figure S2A**). The cell types exhibit clear distinctions, like the previous report [2] . Next, we increased the minimum distance from 0.5 to 0.75, which led to XCL+ NKs and Cycling T cells being moved closer and a shift in their bordering cell types (**Figure S2B**). Further lowering the number of neighbors from 15 to 8 and spread from 1 to 0.3 flipped megakaryocytes (MKs) inward and next to Memory B cells (**Figure S2C**). Lastly, we decreased repulsion from 1 to 0.5. This led to Cycling T and XCL+ NK cell movement away from one another (**Figure S2D**). These changes demonstrate that the relative position of PBMC cell types in a UMAP is affected by UMAP parameters, but no huge movement is observed for this dataset.

**Figure S2. UMAPs for 35,304 peripheral blood mononuclear cells (PBMCs) from COVID-19 patients.** A) UMAP output using default shinyUMAP parameters. B) UMAP output after increasing min_dist parameter to 0.75. C) UMAP output with parameters from B and lowering n_neighbors to 8 along with spread to 0.3. D) UMAP output after further increasing repulsion to 0.5. In all UMAPs, cells are colored by cell types annotated by the original authors, with megakaryocytes (MKs), XCL positive (+) Natural Killer cells (NKs), and Cycling T cells.

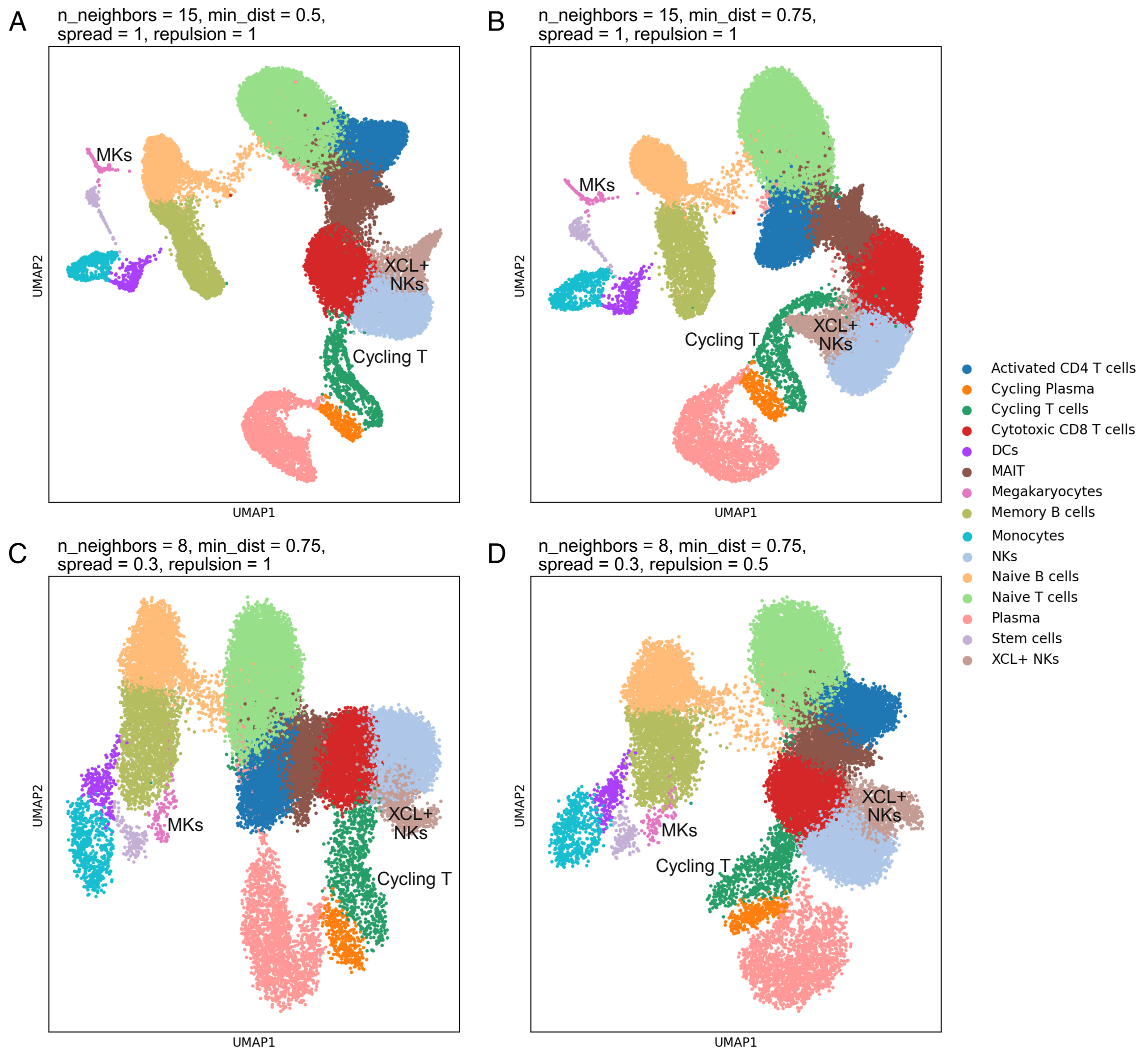


### II. Applying shinyUMAP to a multiomics dataset from PBMCs of healthy individual

The scMultiomics data (scRNA-seq/scATAC-seq) were obtained from 10X Genomics, which was collected from 11,909 PBMCs (referred as “10K”). The integration of the scRNA and scATAC data, cell clustering, and cell type annotation were described in our previously study [4]. The scRNA and scATAC data were uploaded to shinyUMAP separately to generate UMAPs with various parameters (**Figure S3**). The relative positions of cell types in the RNA-based UMAPs and ATAC-based UMAPs show some distinctions. The three cell types, CD4+ naïve, CD4+ memory, and CD8+ naïve T cells, appear to be in a triangle position in the ATAC-based UMAPs, but CD8+ naïve are between CD4+ naïve and CD4+ memory in the

RNA-based UMAPs. The latter seems inconsistent with T cell biology. A comparison of the ATAC-based UMAPs with different parameters (**Figure S3A,C,E**) shows that reducing n_neighbors from 15 to 5 causes the T cell populations to spread apart, with CD4+ memory T cells shifting away from the CD4+ naive/CD8+ naive supercluster. Further decreasing min_distance to 0.1 results in tighter, more defined clusters, particularly compressing the monocyte-mDC continuum while maintaining their linear arrangement. A comparison of the RNA-based UMAPs (**Figure S3B,D,F**) shows that lowering n_neighbors to 5 causes the monocyte populations to elongate and separate from each other, breaking their continuous gradient seen in the default parameters. Further reducing min_distance to 0.1 creates extremely tight clusters with large gaps between cell types, particularly isolating the pDCs and creating artificial separation between related myeloid populations. Comparing ATAC and RNA modalities from the same cells reveals striking differences in UMAP stability. RNA-based UMAPs maintain biologically coherent relationships across parameter changes, better preserving the monocyte-DC developmental continuum as a connected gradient. In contrast, ATAC-based UMAPs fragment these same continuities into discrete populations, with related cell types becoming separated. These differences highlight how UMAPs across different modalities may create misleading impressions of cellular discreteness that may obscure or exaggerate real biological transitions.

**Figure S3. UMAPs of 10K PBMC scMultiomics data.** A) UMAP of the scATAC-seq data using default shinyUMAP parameters. A) UMAP of the scRNA-seq data using default shinyUMAP parameters. C) UMAP of the scATAC-seq data after changing n_neighbour to 5. D) UMAP of the scRNA-seq data after changing n_neighbors to 5. E) UMAP of the scATAC-seq data after changing n_neighbour to 5 and min_distance to 0.1. F) UMAP of the scRNA-seq data after changing n_neighbors to 5 and min_distance to 0.1.

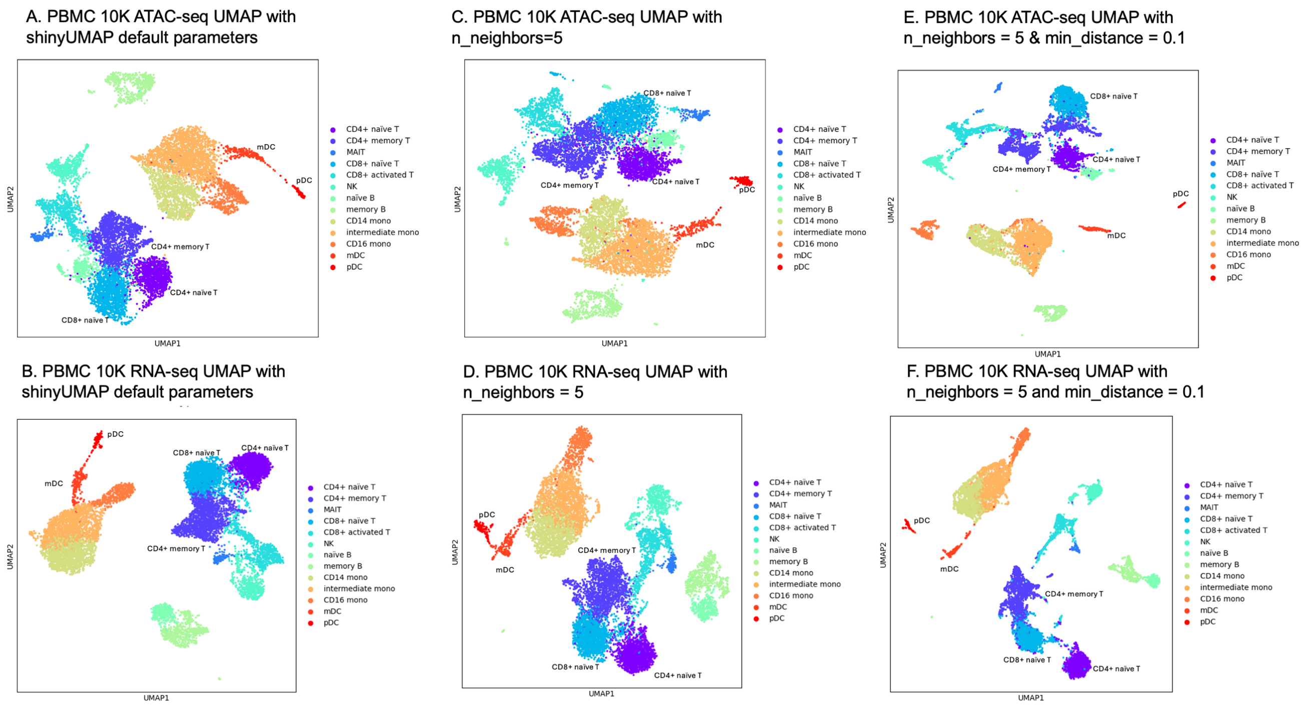


### III. Applying shinyUMAP to mouse embryonic heart development scRNA-seq data

The scRNA-seq data were collected for mesodermal cell linages (*Mesp1*$^{Cre}$) from the mouse pharyngeal apparatus and heart at embryonic stage 9.5 (E9.5), as described previously [5,6]. The data were obtained from both wild type (*WT*) and lineage specific *Tbx1* deletion hearts (*cKO*) for studying the *Tbx1*'s roles in cardiopharyngeal mesoderm development. We extracted the mesodermal cells, containing 23,781 cells in 15 clusters, based on the authors' original analysis [6] . We uploaded the authors' processed data and generated a UMAP using the default settings in shinyUMAP (**Figure S4A**) and then adjusted a few parameters to examine how the UMAPs were affected (**Figure S4B-D**). When the resulting UMAPs are compared, the relative distances of a few clusters are clearly changed. Cluster 10 is close to cluster 2, 4, and 14 using default parameters (**Figure S4A**), but the distances between cluster 10, 2, and 14 become larger when the minimal distance is changed from 0.5 to 0.1 (**Figure S4B**). Cluster 14 moves further away from cluster 10 when the repulsion strength is increased from 1 to 2 (**Figure S4D**). Interestingly, cluster 15 is shown as a “bridge” between cluster 5 and cluster 9 in the default UMAP setting (**Figure S4A**), but its “connection” to cluster 9 disappears when the parameters are changed (**Figure S4B-D**). The latter is more consistent with heart development biology.

**Figure S4. Comparison of UMAPs for mouse embryonic mesodermal scRNA-seq data.** A) UMAP from default shinyUMAP parameters. B) UMAP after changing min_distance to 0.1. C) UMAP after further changing n_neighbors to 30 and spread to 3. D) UMAP after additional change of repulsion to 2. C1 to C15 are, C1: Lung PC, lung progenitors; C2: ST, septum transversum; C3: CT, connective tissues; C4: Sk/L, skeleton/limb; C5: CMs, cardiomyocyte progenitor cells; C6: pSHF, posterior CPM; C7: MLP, multilineage progenitors; C8: BrM, branchiomeric muscle progenitors; C9: aSHF/SoM, anterior CPM and somatic mesoderm; C10: pSHF, posterior CPM; C11: PEO, proepicardial organ; C12: ST, septum transversum; C13: MCs, mesenchyme cells; C14: Sk/L, skeleton/limb; C15: OFT CM, cardiomyocyte progenitor cells in OFT.

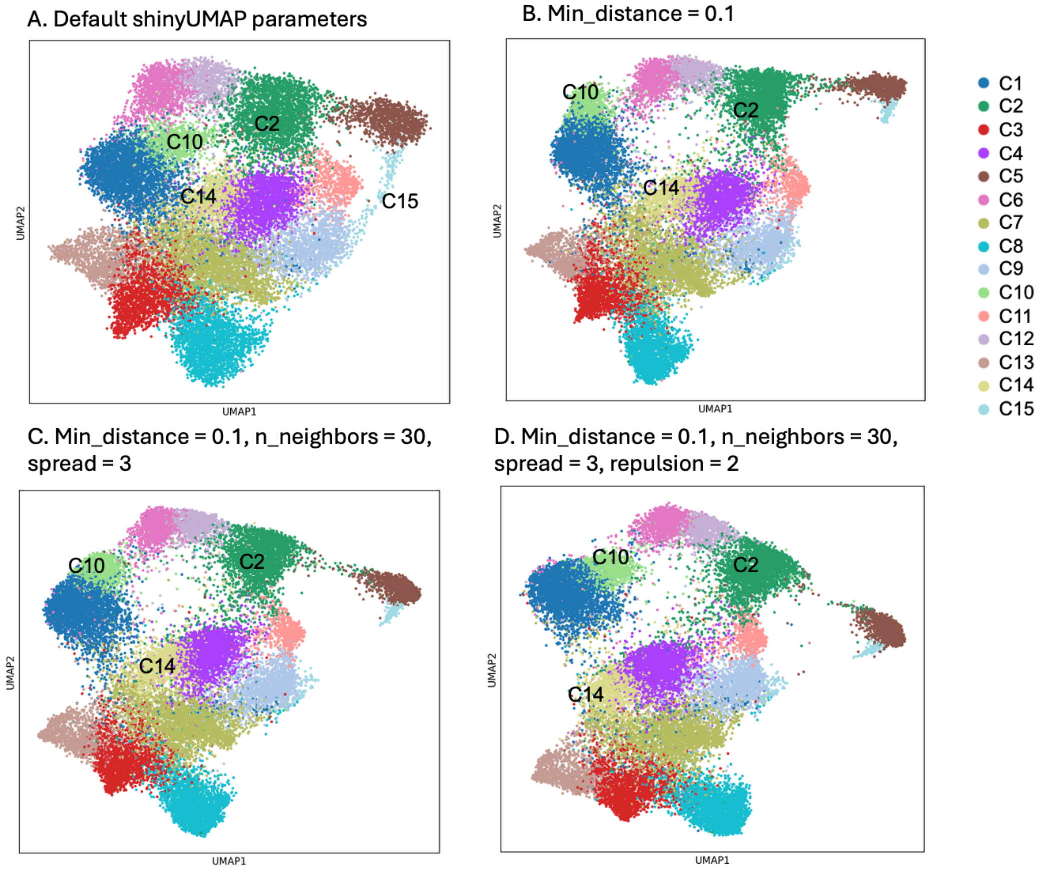

**IV. Applying shinyUMAP to human cortical organoid scRNA-seq datasets.**

Single cell RNA-seq data for 42,187 cells were collected from patient-derived cortical organoids from a previous study, which investigated the effects of 22q11.2 haploinsufficiency on early brain development [7]. We reprocessed the scRNA-seq data using scDAPP software [8] to generate clusters and annotate the cell types using markers from the original study, because the authors did not share their clustering information. The Seurat [9] data object was converted to Anndata [10] format and filtered for top 3000 highly variable genes. The input Anndata object was further downsampled to 10,000 cells using shinyUMAP before computing UMAPs. We first generated an UMAP using the default shinyUMAP setting (**Figure S5A**), then adjusted several parameters to document changes. By increasing n_neighbors from 15 to 30 and decreasing min_distance from 0.5 to 0.1, the relative distance of radial glia and choroid plexus cells increased slightly, in addition to a flip of cluster positions in the UMAP1 (**Figure S5B**). Similarly, when comparing different min_distance under increased n_neighbors, spread and repulsion_strength conditions, the relative distance between radial glia and choroid plexus increased with decreased min_distance (**Figure S5C,D**). However, the relative positions among the cell types in this dataset were not affected significantly by the UMAP parameters. Interestingly, the few actively cycling cells are always with the radial glia, which are prognitors that can divide.

**Figure S5. Shinyumap outputs using human cortical organoid data of 22q11.2 deletion syndrome patients and healthy controls.** A) UMAP with default parameters. B) UMAP after increasing n_neighbors from 15 to 30, and decreasing min_distance from 0.5 to 0.1. C) UMAP with n_neighbors increased from 15 to 30, min_distance increased from 0.5 to 1, spread from 1 to 3, and repulsion strength from 1 to 2. D) UMAP with parameters from C but decreased min_distance from 0.5 to 0.1.

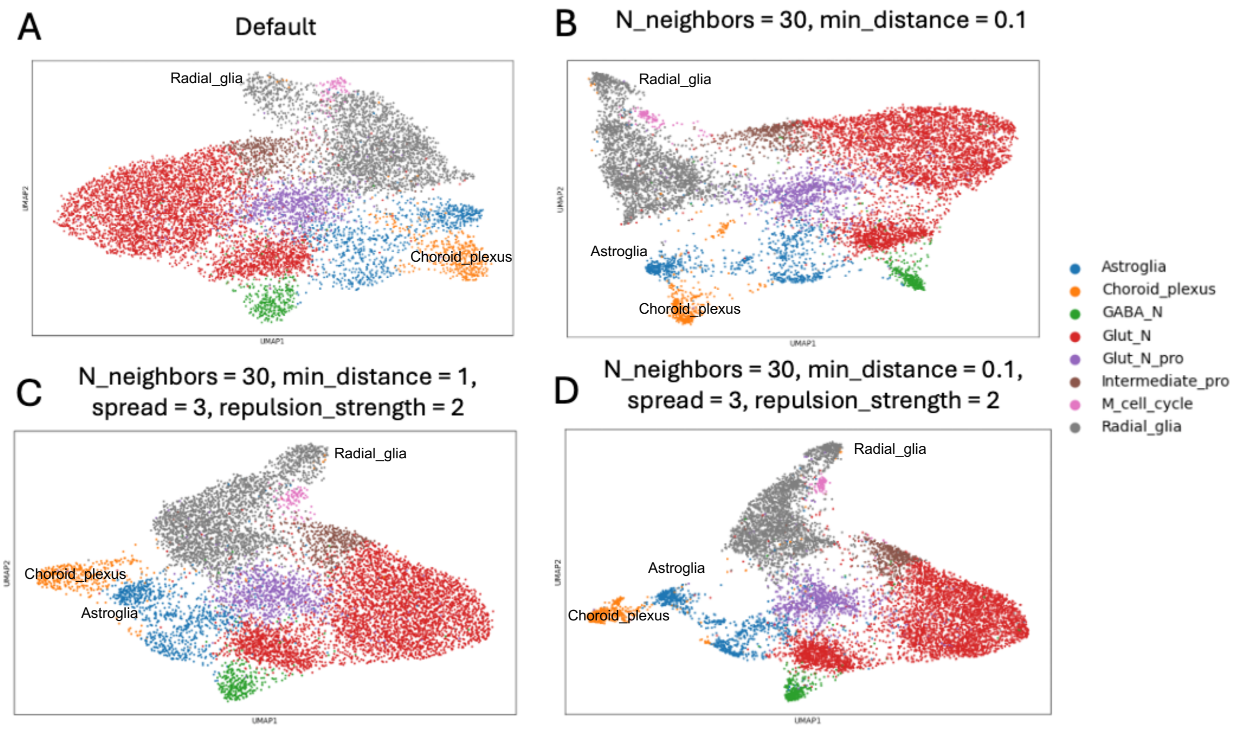
